# An oligo walk to identify siRNAs against the circular Tau 12->7 RNA

**DOI:** 10.1101/2025.01.27.635119

**Authors:** Justin R. Welden, Giorgi Margvelani, Megan Miaro, David Mathews, David W. Rodgers, Stefan Stamm

## Abstract

Circular RNAs are associated with numerous diseases and recent evidence shows that they can be translated into proteins after undergoing RNA modification. Circular RNAs differ from their ‘linear’ mRNA counterparts in their backsplice site, allowing selective targeting using RNA interference, which however limits the options to place the siRNA. We tested all possible siRNAs against the backsplice site of the circTau 12->7 RNA after it was subjected to adenosine to inosine RNA editing, a modification that promotes translation of the circRNA. Most siRNAs reduced the circRNA and protein abundance, which however did not correlate. We identified an siRNA with an IC50 of 750 pmol efficacy on protein expression. This circRNA fulfilled six of the eight criteria for siRNAs targeting mRNAs. Thus, modified circRNAs expressing protein can be targeted with siRNAs, but their optimal sequence needs to be determined empirically.

## Introduction

### Circular RNAs

Circular RNAs (circRNAs) are covalently linked RNAs that in humans are mostly generated by backsplicing, where a 5’ splice site is connected to an upstream 3’ splice site, creating an RNA circle [1]. Although their expression has been correlated with numerous diseases [2], the function of most circRNAs is unclear. An increasing number of circRNAs were shown to be translated [3]. Frequently, the protein product of the circRNAs is directly linked to the disease, suggesting that circRNA removal could be a therapeutic strategy. RNA interference using siRNAs and shRNAs has been used to target the backsplice site that is unique for the circRNA [4]. Although using the backsplice site generates selectivity, it limits the options for siRNA placement.

### siRNAs

siRNAs are 21-23 nt long dsRNAs with a two nucleotide-long 3’ overhang. The dsRNAs are incorporated into the RNA silencing complex, where one strand is loaded on argonaute proteins and guided by this RNA strand, the RNA-argonaute complex cleaves target RNAs [5]. Similarly, shRNAs containing a paired stem and connecting loop can be processed, loaded on argonaute proteins and used to target circRNAs [4, 6]. Several tools have been devised to find the optimal siRNA target sequence [7]. These algorithms take thermodynamic stability, folding free energy change released during siRNA binding, flanking regions, and possible RNA structures into account [8]. An early screen for effective siRNAs showed eight criteria: (i) (30–52% G/C content); (ii) at least three or more A/U nucleotides at the 3’-terminus of the sense strand; (iii, iv) an A at position 19 and 3, (v), a U at position 10, and (vi) no G or C at position 19 and (vii) a G at position 13 and (viii) no internal repeats [9].

### circRNAs generated from the MAPT gene encode proteins

We recently showed that the microtubule associated protein tau generates two circTau RNAs by backsplicing exon 12 to either exon 7 or 10 [10]. Since circRNAs lack a 7-methyl guanosine cap, they rely on internal ribosomal entry sites, which can be provided by base modification. Most circRNAs investigated are translated after methylation of adenosine to N6-methyl adenosine. However, the circTau RNAs are translated after adenosine is modified to inosine [11], with m6A modifications (N6-methyl adenosine) providing a slight additional effect [12]. It has not been systematically studied how siRNAs act on modified RNAs that serve as templates for translation. The circTau 12->7 encoded protein promotes tau aggregation *in vitro* [11], which is a hallmark of Alzheimer’s disease.

To identify possible therapeutic reagents acting on the circTau 12->7 RNA, we tested all possible 20 siRNAs against the A>I edited circTau 12->7 backsplice site and monitored the effect on both circRNA and protein expression. We found no correlation between reduction in protein and circRNA expression levels. The siRNA with best efficacy did not confirm to siRNA rules for mRNAs, suggesting that siRNAs against modified circRNAs need to be determined empirically.

## Material and Methods

### siRNA design

siRNAs corresponded to 21 nt of the sense strand and a dTdT overhang, sequences are shown in **Supplemental Figure 1**.

### Cell culture and transfection

Transient transfections were performed in human embryonic kidney (HEK) 293T cells. One µg a of plasmid DNA was mixed with 250 µl of sterile Opti-MEM reduced serum medium (Thermofisher, 31985070). Co-transfections were at a 1:1 ratio (1 µg:1 µg). siRNAs were added to DNA mix at indicated concentrations. 5 µl of Lipofectamine 2000 was mixed with 250 µl Optimem. DNA and Lipofectamine mixes were combined and incubated at room temperature for 20 min. Cell medium was removed, and then DNA Lipofectamine (Life Technologies, 11668030) mix was added to the cells and incubated for 30 mins in 5% CO_2_ at 37° C. Fresh DMEM medium with 10% Fetal Bovine Serum was added directly to the cells for a total volume of 2 mls after the 30 min incubation. The cells were cultured in a 6-well dish (VWR, 10062-892) and transfected at 60% confluency. The cells were lysed and analyzed 96 h post-transfection.

### Immunoprecipitation of proteins encoded by circTau 12->7

Transfected cells were washed 3X in cold PBS and then lysed in low RIPA buffer (50 mM Tris-HCl pH 7.8, 150 mM NaCl, 1% NP-40 + Fresh protease cocktail inhibitor). Cells were spun for 10 mins at 14,000g in 4° C. The protein encoded by circTau 12->7 is heat stable. Lysates were boiled at 95° C for 5 mins and spun at 14,000g. The circTau 12->7 encoded protein was isolated through immunoprecipitation from cellular extracts using 10 µl of M2 anti-Flag magnetic beads per 1 mg of total protein.

### Measuring circRNAs

Circular RNAs were measured using a custom made Taqman probe as described [12]. Briefly, we used a one-step TaqMan probe kit (TaqMan^™^ RNA-to C_T_^™^ *1-Step* Kit, Invitrogen). The Taqman probe 12->7 was FAM–ACCATCAGCCCCCTTTTTATTTCCT-MGBNFQ. For GAPDH loading control, the GAPDH TaqMan Probe, Invitrogen, Hs02786624_g1 was used. Primers used for circtau were 12 to 7 set2 F: TCGAAGATTGGGTCCCTGGA and 12 to 7 set2 R: TTTTGCTGGAATCCTGGTGG.

### Quantifying siRNA inhibition of circTau expression

Inhibition of circTau expression was quantified by transfecting different amounts of siRNA and measuring protein expression by Western blotting with an anti-Tau antibody. Band intensities were integrated with Image J. Integrated intensities were converted to percent values by dividing independent replicates by the replicate average for the number of siRNA control and multiplying by 100. IC50 values were estimated by nonlinear regression in GraphPad Prism (version 10.4.1) using the log[inhibitor] vs. normalized response – variable slope model with least squares fitting and no weighting [13]. siRNA concentrations were expressed as nanomolar values, and a value of 1 x 10^-6^ nM was used for the no siRNA control in order to allow conversion to a log value Results.

## Results

### siRNA design and experimental setup

The MAPT gene generates several circular RNAs, including one made by backsplicing exon 12 to exon 7, circTAU 12->7 (**Figure 1A**). To study the effect of siRNAs on both protein and circRNA expression, we generated reporter genes that contain the cDNA of the circular RNA, flanked by the natural introns (**Fig. 1B**). Upon transfection into HEK293 cells, this construct generates a circRNA, which has one start codon and a 681 nt long open reading frame, with no stop codon (**Fig 1C**) [11]. The circTau 12->7 RNA is translated after undergoing adenosine to inosine modifications, caused by ADAR (adenosine deaminase acting on RNA) enzymes. When inosines are present on the circRNA, translation occurs generating a 40 kDa protein using an unknown termination mechanism. The protein encoded by circTau 12->7 is identical to the microtubule binding repeat region of the mRNA-encoded protein, with only two amino acid difference corresponding to the exon 12 to exon 7 junction. To detect the protein, we thus introduced a 3x Flag tag (**Fig. 1B**). The circTau encoded protein is heat stable and can be recovered by immunoprecipitation after boiling transfected cells in lysis buffer. The amount of protein generated from the circRNA does not correlate with the amount of circRNA but depends on the RNA modification present [12].

**Figure 1:**
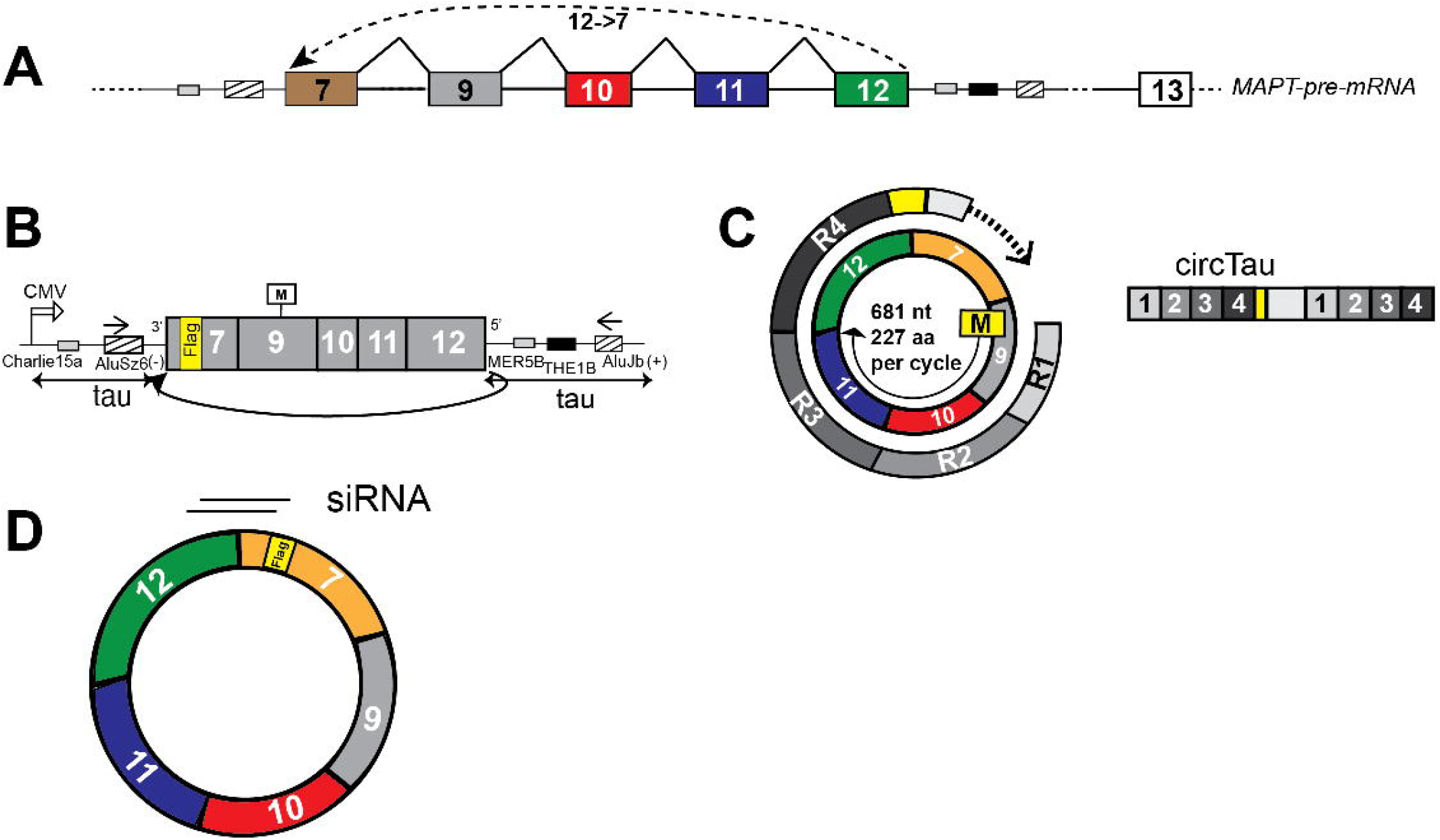
Testing strategy. **A**. Structure of the MAPT gene. Two circular RNAs generated by backsplicing of exon 12 to either exon 7 or exon 10 are indicated. **B**. Expression construct used to generate 12->7 circTau in transfection assays. The introns between Exons 7 to 12 were removed and an in-frame FLAG tag was inserted into exon 7. This cDNA was inserted between the natural introns flanking exon 7 and 12, respectively. **C**. Structure of the 12->7 circRNA that is shown as a circle in the middle. Upon A>I editing, the circTau RNA is translated from a single start codon into a protein that contains microtubule binding repeats 1-4 (R1-R4). The structure of the proteins is shown in the right. **D**. Location of the siRNAs against the 12->7 backsplice site.

Little is known about the effect of siRNAs on circRNAs or heavily modified circRNAs. Because the location of the siRNA is determined by the backsplice site, the area for siRNA selection is limited and no siRNA confers to the recommended design features. We thus performed an siRNA walk, testing all possible siRNAs covering the backsplice junction (**Fig. 1D**). The siRNAs were 21 nt long, plus two dT nucleotides at the 3’ overhangs. The sequences of the siRNAs are given in **Supplemental Figure S1.**

### Effect on RNA

To test all siRNAs, we transfected siRNAs in 100 and 1000 nM concentration together with the circTau 12->7 expression clone, as well as a construct expressing ADAR-p150. From these transfections, one aliquot was used to isolate total RNA and another one was used to test circTau 12->7 encoded protein expression. We determined the amount of circTau RNA using a taqman assay we previously developed, where the taqman probe spans the exon junction (**Supplemental Figure S2**). The taqman signal was normalized to GAPDH, measured by a taqman probe (Hs02786624_g1). Each siRNA was tested in triplicates. To account for possible variations during the transfection, the amount of RNA was normalized to the no siRNA control. All siRNAs reduced the circTau 12->7 RNA expression. However, there are marked differences, with siRNA 7_15 having the lowest effect and siRNA 7_2, 15 having the strongest effect at 100 nM concentration. To test different expression constructs, we also tested a circTau 7->12 expression construct that had the Flag-tag removed near the backsplice site, which did not change overall siRNA efficacy (**Table 1, Supplemental Figure 3A, B**).

**Table 1.**
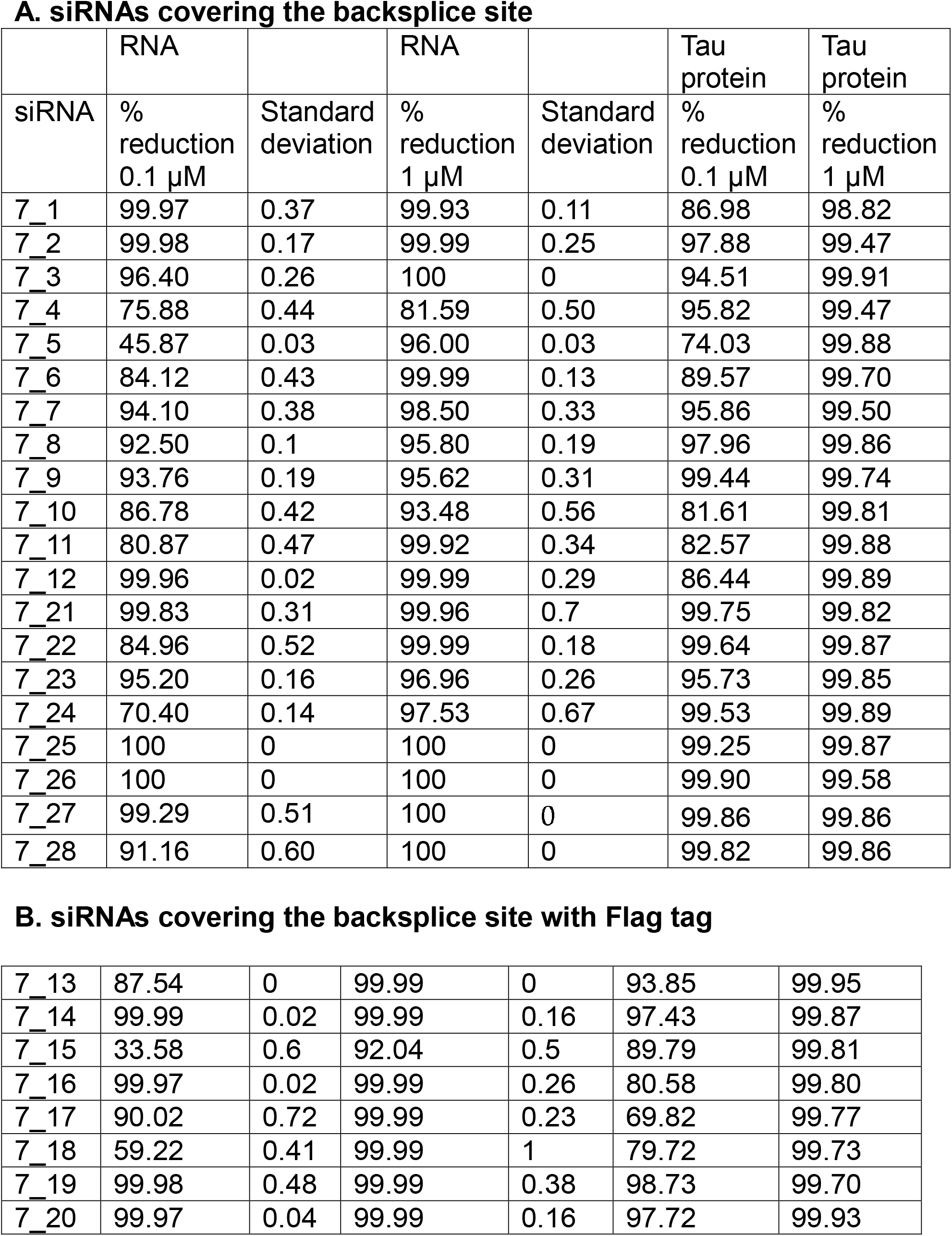
Sequence of siRNAs. The partial sequence of exons 12 and 7 of the backsplice site is indicated. The sequences of the sense (passenger strand) of the siRNAs are shown. All siRNAs contain dT overhangs. As an example, the 7-1 siRNA duplex is shown below.

|  | RNA |  | RNA |  | Tau protein | Tau protein |
| --- | --- | --- | --- | --- | --- | --- |
| siRNA | % reduction<br>0.1 μM | Standard deviation | % reduction<br>1 μM | Standard deviation | % reduction<br>0.1 μM | % reduction<br>1 μM |
| 7_1 | 99.97 | 0.37 | 99.93 | 0.11 | 86.98 | 98.82 |
| 7_2 | 99.98 | 0.17 | 99.99 | 0.25 | 97.88 | 99.47 |
| 7_3 | 96.40 | 0.26 | 100 | 0 | 94.51 | 99.91 |
| 7_4 | 75.88 | 0.44 | 81.59 | 0.50 | 95.82 | 99.47 |
| 7_5 | 45.87 | 0.03 | 96.00 | 0.03 | 74.03 | 99.88 |
| 7_6 | 84.12 | 0.43 | 99.99 | 0.13 | 89.57 | 99.70 |
| 7_7 | 94.10 | 0.38 | 98.50 | 0.33 | 95.86 | 99.50 |
| 7_8 | 92.50 | 0.1 | 95.80 | 0.19 | 97.96 | 99.86 |
| 7_9 | 93.76 | 0.19 | 95.62 | 0.31 | 99.44 | 99.74 |
| 7_10 | 86.78 | 0.42 | 93.48 | 0.56 | 81.61 | 99.81 |
| 7_11 | 80.87 | 0.47 | 99.92 | 0.34 | 82.57 | 99.88 |
| 7_12 | 99.96 | 0.02 | 99.99 | 0.29 | 86.44 | 99.89 |
| 7_21 | 99.83 | 0.31 | 99.96 | 0.7 | 99.75 | 99.82 |
| 7_22 | 84.96 | 0.52 | 99.99 | 0.18 | 99.64 | 99.87 |
| 7_23 | 95.20 | 0.16 | 96.96 | 0.26 | 95.73 | 99.85 |
| 7_24 | 70.40 | 0.14 | 97.53 | 0.67 | 99.53 | 99.89 |
| 7_25 | 100 | 0 | 100 | 0 | 99.25 | 99.87 |
| 7_26 | 100 | 0 | 100 | 0 | 99.90 | 99.58 |
| 7_27 | 99.29 | 0.51 | 100 | 0 | 99.86 | 99.86 |
| 7_28 | 91.16 | 0.60 | 100 | 0 | 99.82 | 99.86 |
| **B. siRNAs covering the backsplice site with Flag tag** |  |  |  |  |  |  |
| 7_13 | 87.54 | 0 | 99.99 | 0 | 93.85 | 99.95 |
| 7_14 | 99.99 | 0.02 | 99.99 | 0.16 | 97.43 | 99.87 |
| 7_15 | 33.58 | 0.6 | 92.04 | 0.5 | 89.79 | 99.81 |
| 7_16 | 99.97 | 0.02 | 99.99 | 0.26 | 80.58 | 99.80 |
| 7_17 | 90.02 | 0.72 | 99.99 | 0.23 | 69.82 | 99.77 |
| 7_18 | 59.22 | 0.41 | 99.99 | 1 | 79.72 | 99.73 |
| 7_19 | 99.98 | 0.48 | 99.99 | 0.38 | 98.73 | 99.70 |
| 7_20 | 99.97 | 0.04 | 99.99 | 0.16 | 97.72 | 99.93 |

### Effect on Protein

To detect the effect of siRNAs on circTau protein formation, we measured proteins made in the presence of siRNAs. The protein was generated from the circTau 12->7 expression construct in the presence of transfected ADAR1 that caused adenosine to inosine edition of the circRNA. We immunoprecipitated the heat stable circTau protein from boiled lysates using the Flag tag. The circTau protein in the immunoprecipitates was detected using a flag and anti tau antibody. For each circRNA, we next normalized the protein signal to the control siRNA and found the strongest effect for siRNA 7_26 (**Table 1, Supplemental Fig. 4**).

### No Effect on linear MAPT

To test the selectivity of the siRNA for circular RNAs, we tested the siRNA sequences with the highest efficacy against the mRNA encoded 0N4R tau isoform, which was expressed from a cDNA construct. For siRNA 7-8, we did not see a significant effect on either RNA or protein expression of mRNA encoded MAPT (**Figure 2A**). This siRNA exhibits complementarity to 13 and 8 nt in exon 12 and 7, respectively. We further determined its overall efficacy on circTau 12->7 RNA-encoded protein, using an siRNA range from 1 nM to 1 µM. We found an IC50 of 170 pM 95% CI [90 to 280 pM], which is comparable to other siRNAs [14] (**Figure 2B, C**).

**Figure 2:**
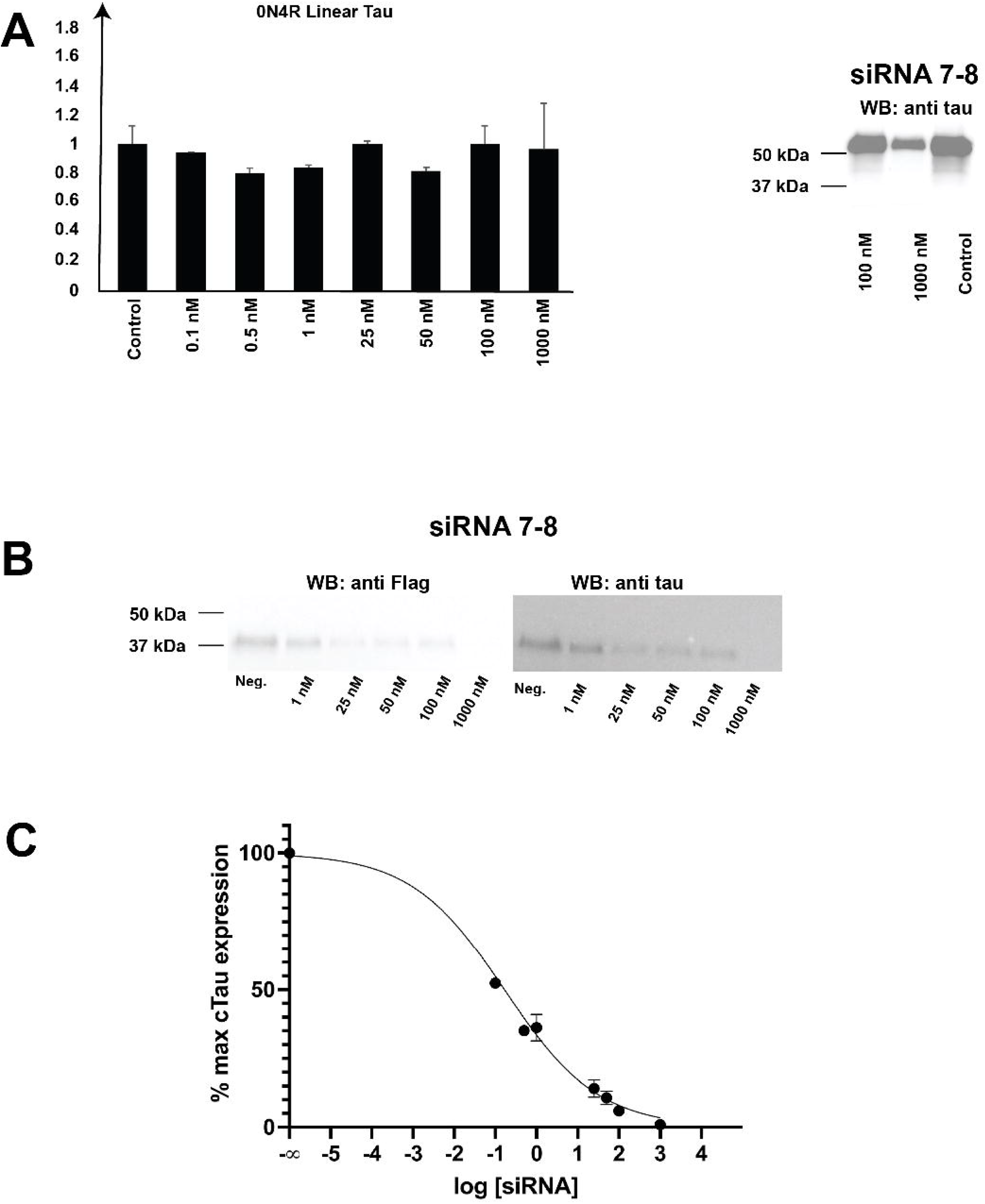
Quantification of siRNA7-8. **A**. Left: An expression clone for tau N04R was cotransfected with siRNA 7-8 at the concentration indicated. The concentration of 0N4R mRNA was determined using a taqman probe (Thermofisher, 4331182). Right: circTau 12->7 protein was analyzed by Western blot. **B**. The circTau 12->7 expression construct was cotransfected with various concentration of the siRNA7-8 and protein was detected by immunoprecipitation followed by Western blot. **C**. Dose-response curve for siRNA inhibition of circTau expression. A plot of circTau expression quantified by anti-Tau antibody detection on a Western blot versus siRNA 7-8 concentration is shown. Values for circTau expression are given as the percentage of circTau expression for the no siRNA control and are plotted against the log of the nanomolar siRNA concentration. Error bars indicate the standard error of the mean for independent replicates. The IC50 is 170 pM with 95% CI [90 to 280 pM].

## Discussion

### siRNAs can be used to target edited circRNAs

Here, we tested whether siRNAs can interact with A>I edited circRNAs targeting the unique backsplice site. In the presence of ADAR1, most adenosines of the circRNA are changed into inosines at low levels [11]. Thus, the circRNAs likely represent a mixture of modified molecules. We found that these widespread, low-level modifications do not prohibit siRNA mediated circRNA decay.

### No correlation between protein and RNA effect

We previously found that the levels of circRNA expression do not correlate with the amount of encoded proteins [12]. The siRNA data in Table 1 support this observation, indicating that the amount of protein made from circRNAs does not correlate with the amount of RNA.

### Determinants of siRNA efficacy

Our best siRNA had IC50 of 170 pM for protein, which is comparable with other siRNAs. For mRNAs, rules for siRNAs have been described that are the basis of bioinformatic prediction programs [9, 15]. Since we target the backsplice site, we could not choose the placement for the siRNA. Despite these limitations, we identified numerous siRNA that reduce the circTau12->7 RNA. None of these active siRNAs confirm to the eight rules for siRNA design [9]. Supplemental Figure 6 shows the adherence of the siRNAs with the eight design rules. Noticeably, our best siRNA, siRNA 7-8, fulfilled five of the eight conditions.

In summary, siRNAs against circular RNA backsplice sites adhere to rules determined for mRNAs when the RNA is considered. However, their effect on protein production could not be predicted, which could reflect the modification dependency of siRNA translation. Thus, siRNAs aimed to reduce proteins made from circular RNAs should be designed empirically.

## Supplemental Figures

***Supplemental Figure S1: siRNA sequences***

**A**. The backsplice site of exon 12 to exon 7 of the MAPT gene is indicated by different shading. siRNAs used are shown underneath.

**B**. Example of an siRNA duplex with the uniform dTdT overhangs at the 3’ end.

***Supplemental Figure S2A: Taqman Assay to measure circTau abundance***

Sequence and location of the TaqMan probe spanning exons 12 and 7. Star: FAM, circle: MGBNFQ quencher. The location of the primers in the circRNA is schematically shown on right.

***Supplemental Figure S3 Effect of siRNAs on circTau RNA expression levels***.

1 µg of the circTau expression construct and 1 µg ADAR1-p150 were transfected into HEK293 cells in the presence of siRNAs indicated at 0.1 µM and 1 µM concentrations. The circTau RNA was lysed 4 days post-transfection and the concentration was determined by qPCR, normalized to GAPDH.

***Supplemental Figure S4: Effect of siRNAs on circTau Protein expression levels***.

1 µg of the circTau expression construct and 1 µg ADAR1-p150 were cotransfected into HEK293 cells in the presence of siRNAs indicated at 0.1 µM and 1 µM concentrations. Cells were lysed 4 days post-transfection, and boiled lysates were immunoprecipitated using anti-Flag antibodies. siRNAs targeting the circTau junction drastically reduce protein expression.

