## Supplemental Figures for "An oligo walk to identify siRNAs against the circular Tau 12->7 RNA": 1_Supplemental Figure S1.docx

**Figure S1: Sequences of the siRNAs**

```
A. Sequences

              Exon 12               Exon 7
          CTGGCGGAGGAAATAAAAAGGGGGCTGATGGTAAAACGAA
 7_1      CTGGCGGAGGAAATAAAAAGG
 7_2       TGGCGGAGGAAATAAAAAGGG
 7-3        GGCGGAGGAAATAAAAAGGGG
 7_4         GCGGAGGAAATAAAAAGGGGG
 7_5          CGGAGGAAATAAAAAGGGGGC
 7_6           GGAGGAAATAAAAAGGGGGCT
 7_7            GAGGAAATAAAAAGGGGGCTG
 7_8             AGGAAATAAAAAGGGGGCTGA
 7_9              GGAAATAAAAAGGGGGCTGAT
 7_10              GAAATAAAAAGGGGGCTGATG
 7_11               AAATAAAAAGGGGGCTGATGG
 7_12                AATAAAAAGGGGGCTGATGGT
 7-21                 ATAAAAAGGGGGCTGATGGTA
 7-22                  TAAAAAGGGGGCTGATGGTAA
 7-23                   AAAAAGGGGGCTGATGGTAAA
 7-24                    AAAAGGGGGCTGATGGTAAAA
 7-25                     AAAGGGGGCTGATGGTAAAAC
 7-26                      AAGGGGGCTGATGGTAAAACG
 7-27                       AGGGGGCTGATGGTAAAACGA
 7-28                        GGGGGCTGATGGTAAAACGAA

B. Example of siRNA douplex

7_1.    5’      CTGGCGGAGGAAATAAAAAGGdTdT
        3’  dTdTGACCGCCTCCTTTATTTTTCC


siRNAs for Flag tag
 7_13                 ATAAAAAGGGGGCTGATGGTG
 7_14                  TAAAAAGGGGGCTGATGGTGA
 7_15                   AAAAAGGGGGCTGATGGTGAC
 7_16                    AAAAGGGGGCTGATGGTGACT
 7_17                     AAAGGGGGCTGATGGTGACTA
 7_18                      AAGGGGGCTGATGGTGACTAC
 7_19                       AGGGGGCTGATGGTGACTACA
 7_20                        GGGGGCTGATGGTGACTACAA
```

A. The backsplice site of exon 12 to exon 7 of the MAPT gene is indicated by different shading. siRNAs used are shown underneath.

B. Example of an siRNA douplex with the uniform dTdT overhangs at the 3’ end.
