## Supplementary figures and images for "An oligo walk to identify siRNAs against the circular Tau 12->7 RNA"

### 2_Supplemetal Figure S2A_v2 .png

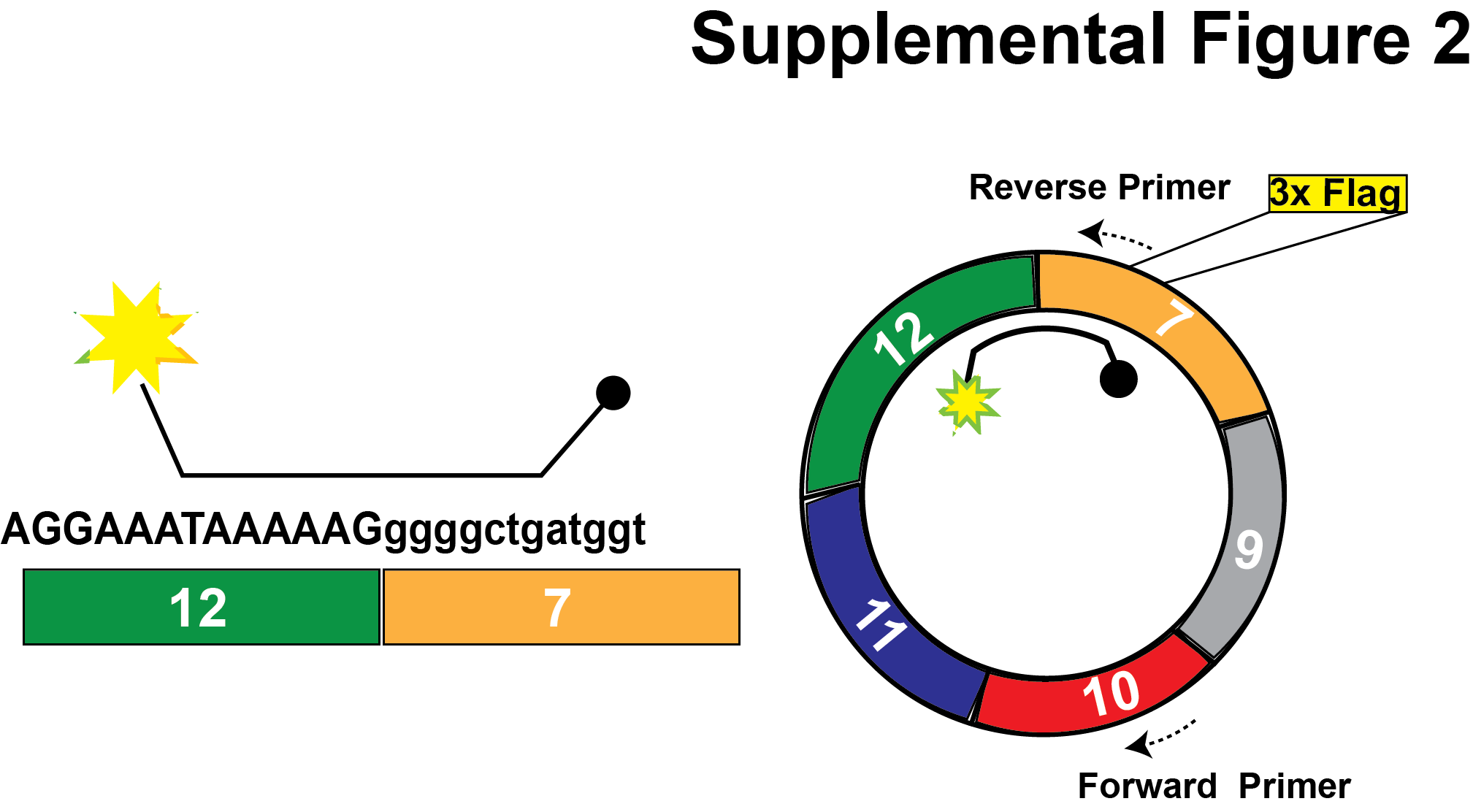

### 3_Supplemental Figure 3A_v4.png

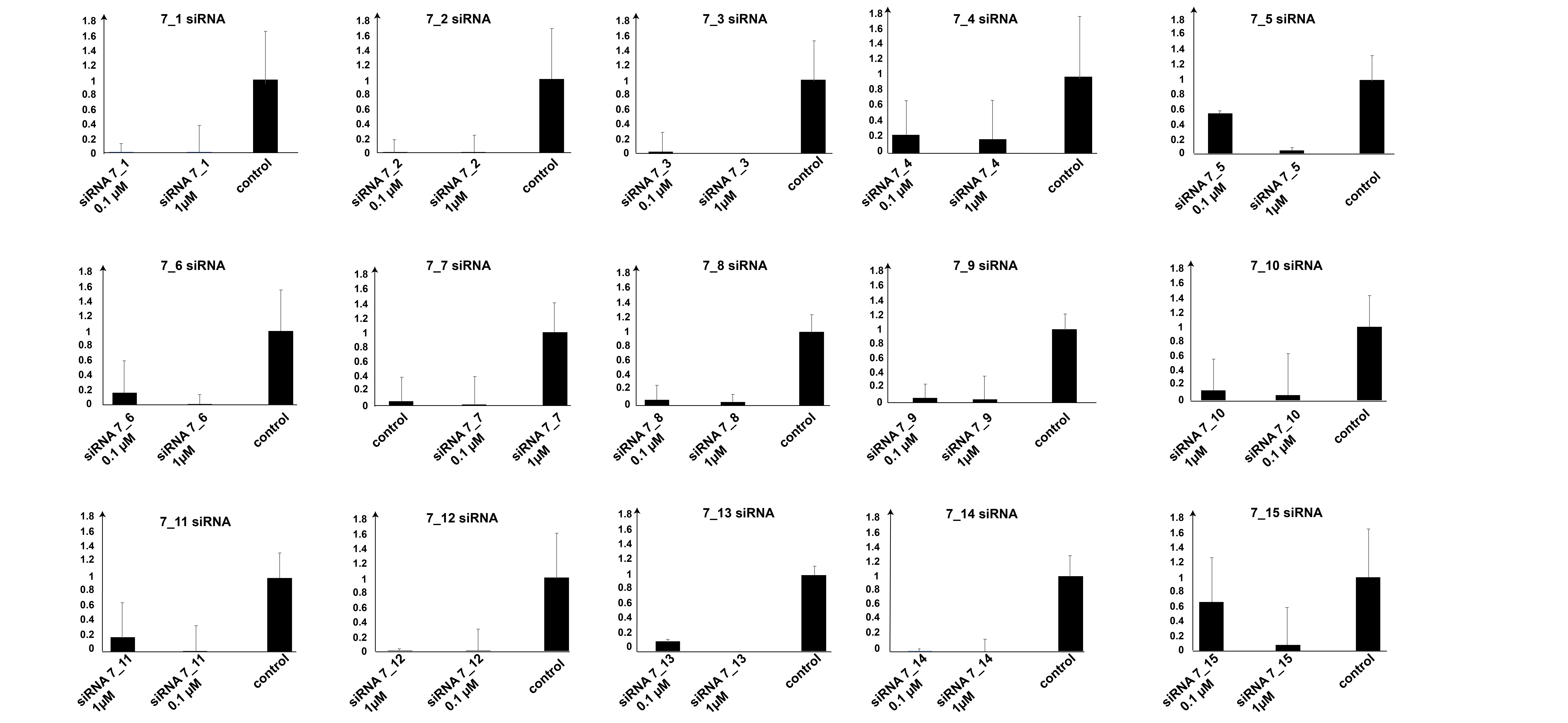

### 3_Supplemental Figure 3B_v4.png

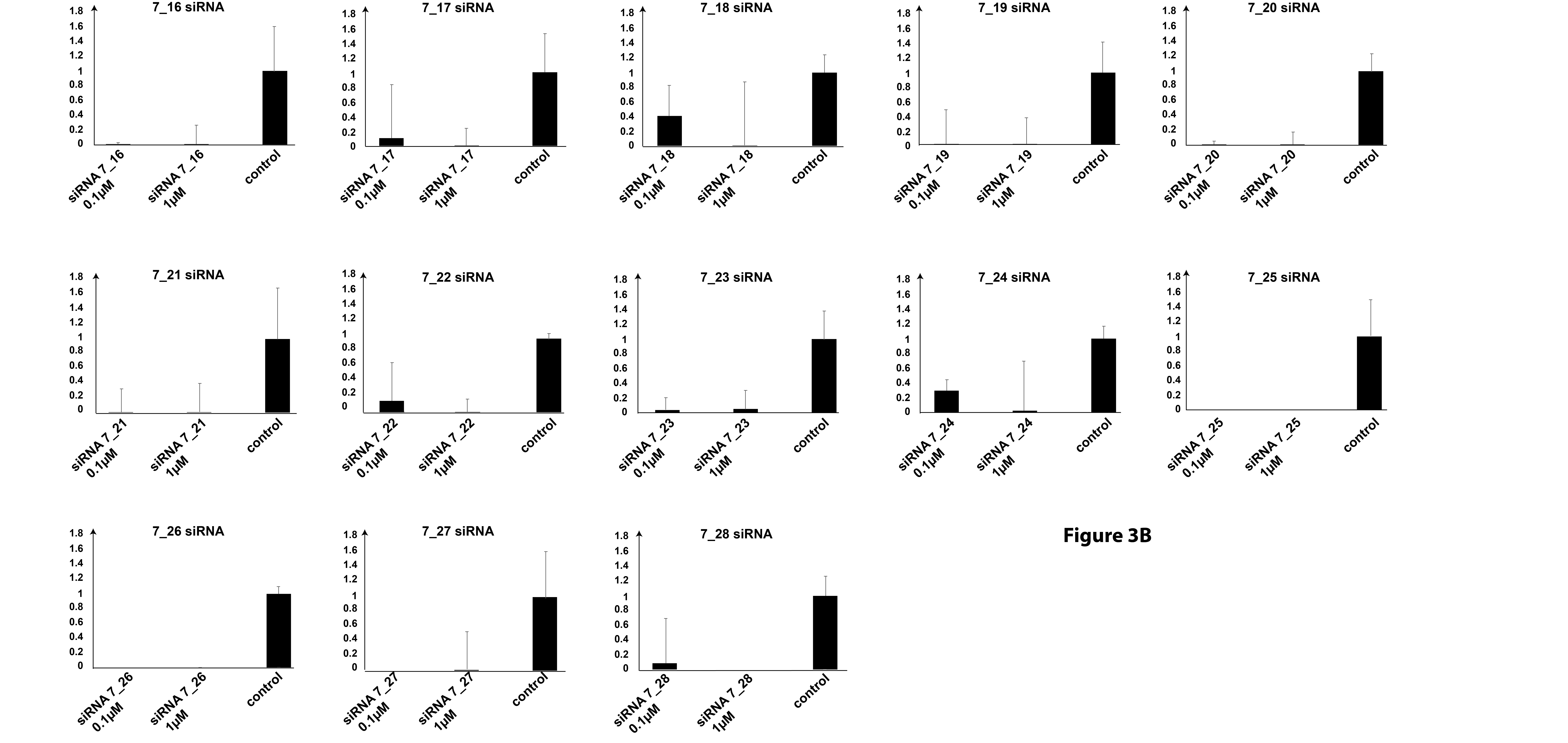

### 3_Supplemental Figure 3D .png

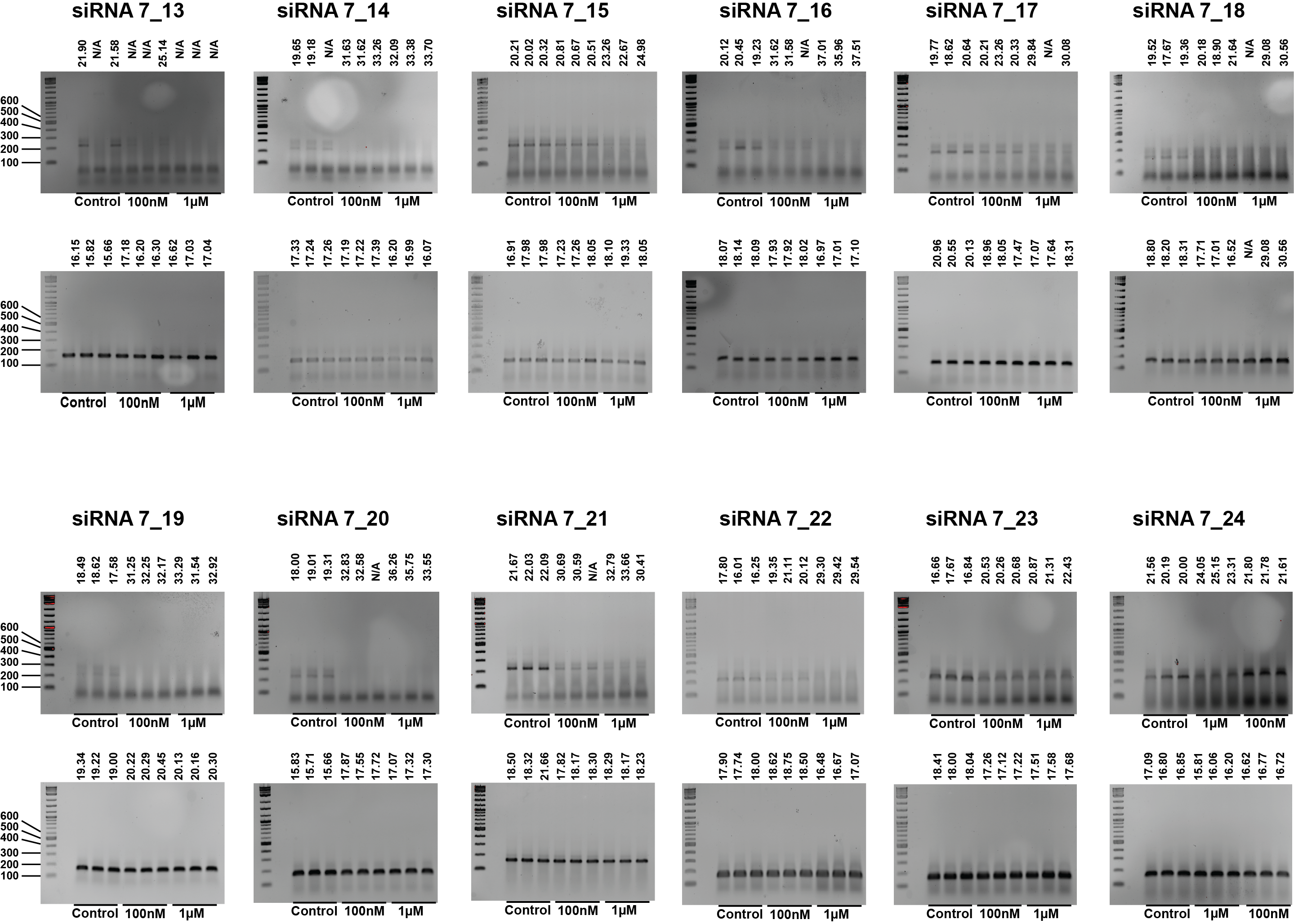

### 3_Supplemental Figure 3E .png

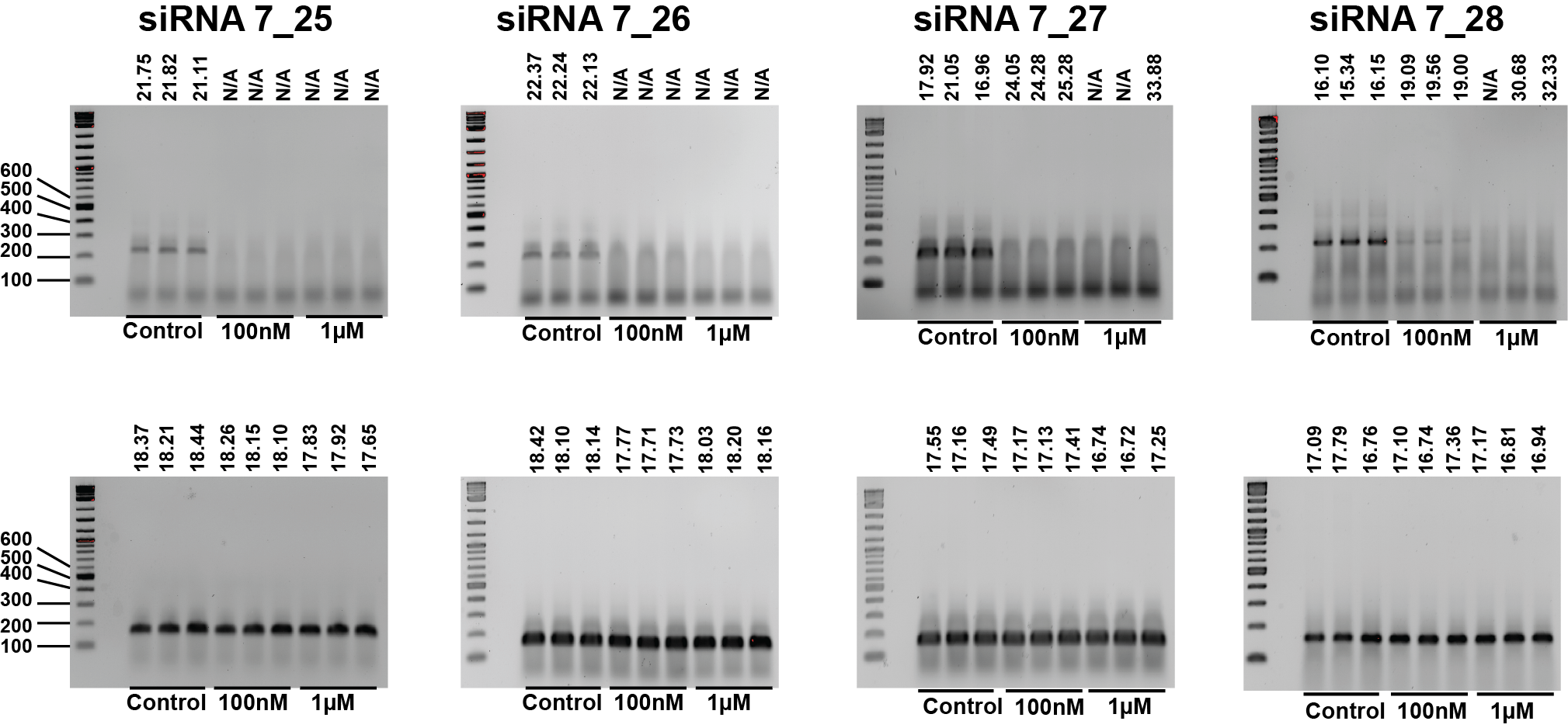

### 4_Supplemental Figure 4_V6.png

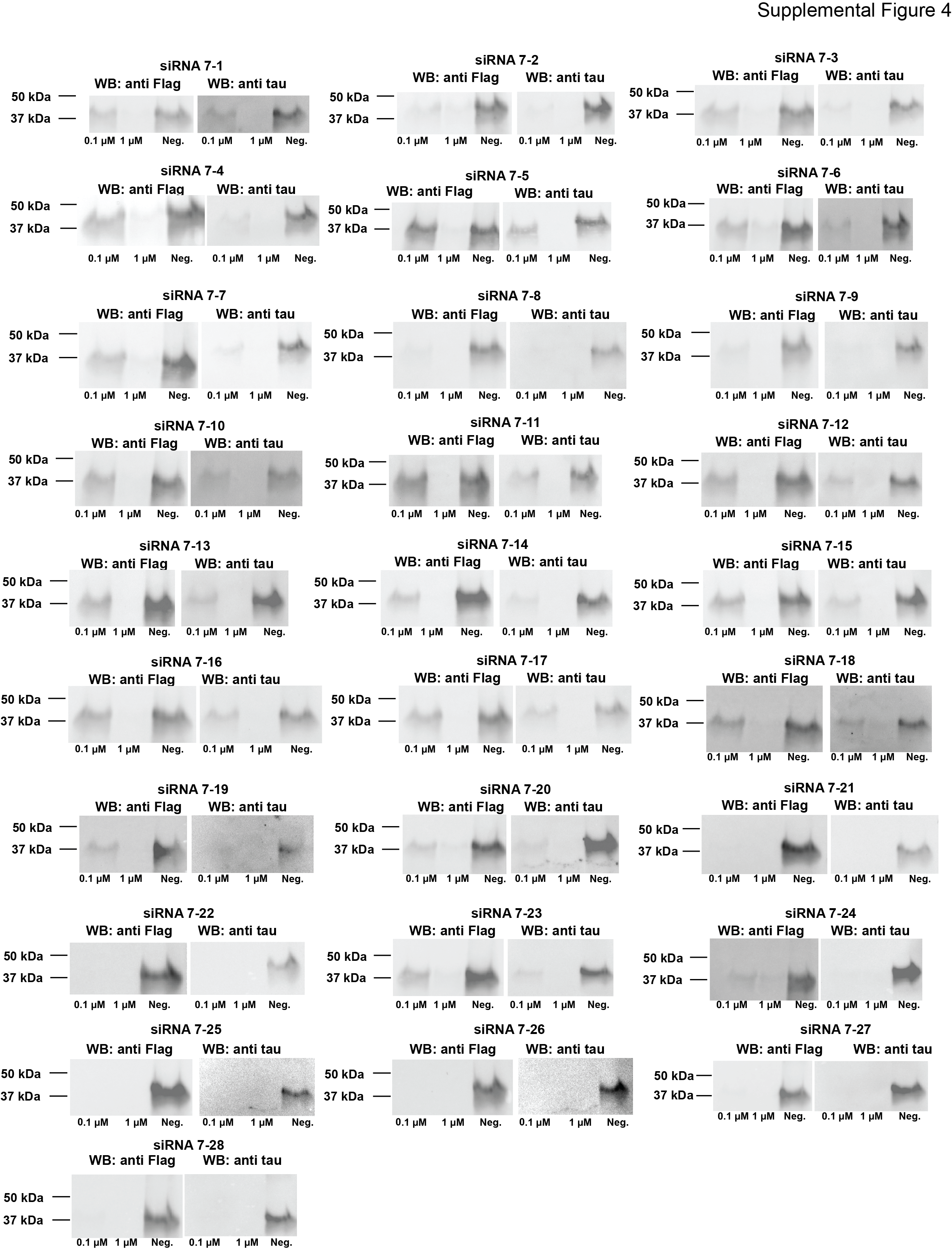

### supplemental figure 3C V2 .png

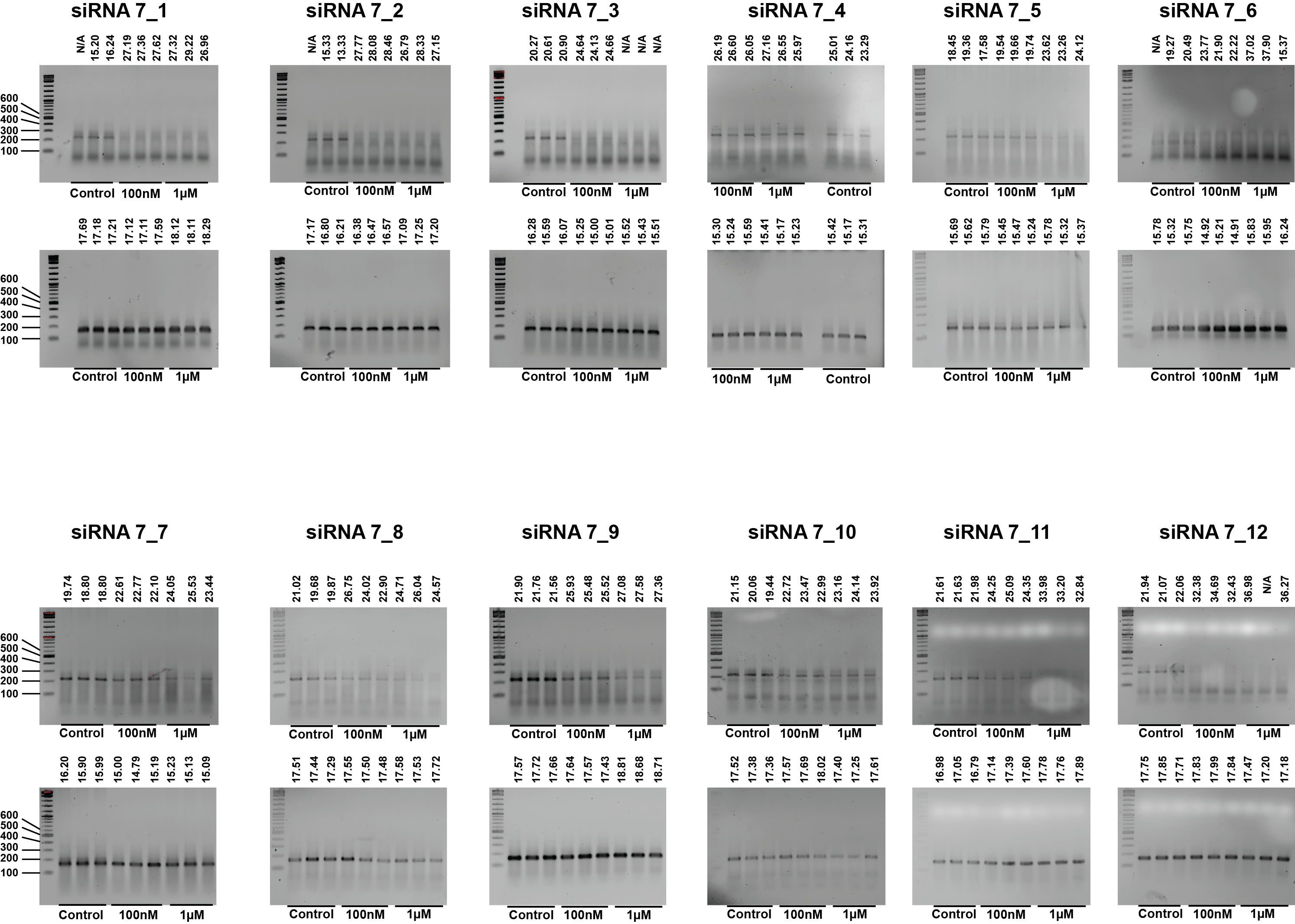
